# AI-Powered Discovery of Novel RNA Viruses from the Permafrost of a 14,300-Year-Old Pleistocene Wolf

**DOI:** 10.64898/2026.09.17.746571

**Authors:** Syed Zaheer ud Din, Qingfa Wu

## Abstract

Ancient viruses preserved as molecular relics offer rare and often unpredictable insights into virus-host co-evolution and the ecological dynamics of past ecosystems. However, the recovery of ancient RNA viruses via paleotranscriptomics has remained largely unexplored, constrained by the inherent chemical instability of RNA and the lack of sensitive detection tools capable of identifying deeply divergent sequences. Here, we leveraged recent advances in artificial intelligence and high-throughput sequencing to conduct comprehensive metatranscriptomic mining of publicly available RNA-seq datasets from three ancient or extinct host species: the woolly mammoth (*Mammuthus primigenius*), the Tasmanian tiger (*Thylacinus cynocephalus*), and the gray wolf (*Canis lupus*). We performed sensitive homology searches using the AI-driven protein language model Lucaprot, coupled with structural validation via AlphaFold2, to screen billions of raw sequencing reads for conserved viral RNA-dependent RNA polymerase (RdRp) signature genes. Our pipeline identified two complete previously unknown RNA viruses in a 14,300-year-old Pleistocene wolf specimen. Phylogenetic analyses placed them within established mycovirus genera (*Duamitovirus* and *Orthocurvulavirus*), indicating they infected fungi that inhabited the carcass rather than the wolf itself. Despite deep sequence divergence from known viruses (57.6% and 60.8% RdRp amino acid identity, respectively), the catalytic A, B, and C motifs remain structurally intact. Strict authentication through multiple analyses firmly verified their ancient provenance. To our knowledge, this is the earliest documented evidence of novel RNA viruses persisting within a host-associated microbiome, extending the observed preservation timescale from centuries to over fourteen millennia. Our findings demonstrate that permafrost is a viable substrate for paleovirological discovery extending beyond the host organism and opening new opportunities for reconstructing ancient microbial and viral ecosystems.

## Introduction

The emergence of high-throughput sequencing has transformed paleogenomics, making it possible to gather and examine genetic material from species that have long been extinct at unparalleled resolution (Danielewski, Żuraszek et al. 2023). The reconstruction of the Neanderthal (*Homo neanderthalensis*) genome, along with other groundbreaking work in ancient DNA sequencing, demonstrated that extraction procedures and powerful computational tools could overcome the hurdles posed by severe molecular degradation and external contamination, revealing insights into deep evolutionary history (Fu, Li et al. 2014). Since then, researchers have used these methods to study extinct megafauna, including the Tasmanian tiger and the woolly mammoth, yielding insights into population dynamics and the ecological causes of extinction (von Seth, Niemann et al. 2018). While ancient genomics has thrived, paleovirology, particularly ancient RNA viruses, is a relatively new field. This distinction stems from a fundamental biological fact that RNA is inherently less stable than DNA, making it particularly challenging to extract from ancient/historical materials. For decades, paleovirological research was primarily focused on endogenous retroviruses, which leave long-lasting molecular fossils integrated into host genomes. Non-retroviral RNA viruses, by contrast, were long thought to leave no identifiable paleontological trace, which makes their prehistoric existence largely enigmatic (Johnson 2019).

Recent developments, however, have started to contest this. Researchers have shown that informative RNA can be obtained from ancient samples preserved in permafrost and from historical skins kept in museum collections (von Seth, Niemann et al. 2018). Moreover, they have recently extracted RNA from a woolly mammoth specimen dated to approximately 40,000–50,000 years ago, pushing the boundaries of ancient RNA recovery into the Pleistocene (Mármol-Sánchez, Fromm et al. 2026). Metatranscriptomic studies of naturally mummified Adélie penguin remains have produced near-complete genomes of known RNA viruses, such as picornaviruses and rotaviruses, from specimens nearly two thousand years old (Hinzke, Lauber et al. 2025). Even in human contexts, alcohol-preserved 18th-century lung tissue has enabled reconstruction of the oldest human-associated virus genome recovered to date (Barnett, Castillo et al. 2026). Together, these findings indicate that RNA preservation, while rare, is not confined to extraordinary situations and can happen under diverse conditions, such as permafrost, drying, ethanol, and formalin treatment. At the same time, large-scale viral metatranscriptomic efforts have revealed that the public sequence read archive (SRA) occasionally contains foreign viral sequences unintentionally sequenced alongside host transcriptomes (Zhu, Raza et al. 2025). These observations and robust pipelines highlight a significant opportunity. Using metatranscriptomic mining on ancient and extinct host transcriptomes could aid in the discovery of RNA viruses that previously infected these creatures or their associated microbiomes. These discoveries would broaden our understanding of viral diversity in past ecological situations and show the evolutionary background of host-virus relationships.

In this study, we hypothesized that transcriptome datasets acquired from tissues of extinct animals, such as the woolly mammoth, Tasmanian tiger, and Pleistocene gray wolf, may contain exogenous RNA viruses that were present in the host or its associated microbiome at the time of death. We developed a multi-tiered computational screening pipeline to examine this (Figure 1). Initially, we ran homology-based searches with Diamond BLASTx against the curated viral RefSeq databases to uncover conserved viral hallmark genes. To capture extremely divergent or extensively altered viral sequences that defy typical homology identification, we also used a protein language model AI tool, Lucaprot, for quick, structure-informed sequence screening (Hou et al. 2024). The candidate RdRp sequences identified were subsequently subjected to tertiary structure prediction using AlphaFold, enabling us to verify the structural conservation of the catalytic palm domain motifs (A, B, and C) independent of primary sequence divergence. This integrated approach enabled us to recover two complete, novel, exogenous RNA viruses from a 14,300-year-old Pleistocene canid. Phylogenetic reconstruction placed these viruses within an established viral clade, yet their extensive sequence divergence supports their classification as putative new species. To our knowledge, this study constitutes the first discovery of exogenous RNA viruses from an ancient Pleistocene canid host, unequivocally demonstrating the feasibility and power of paleotranscriptomic viral discovery, bridging a massive chronological gap in paleovirology, and scaling back the observable timeline of intact animal RNA virus evolution from centuries to over fourteen millennia, demonstrating that biologically active viral sequences can persist within ancient mammalian soft tissues.

**Figure 1.**
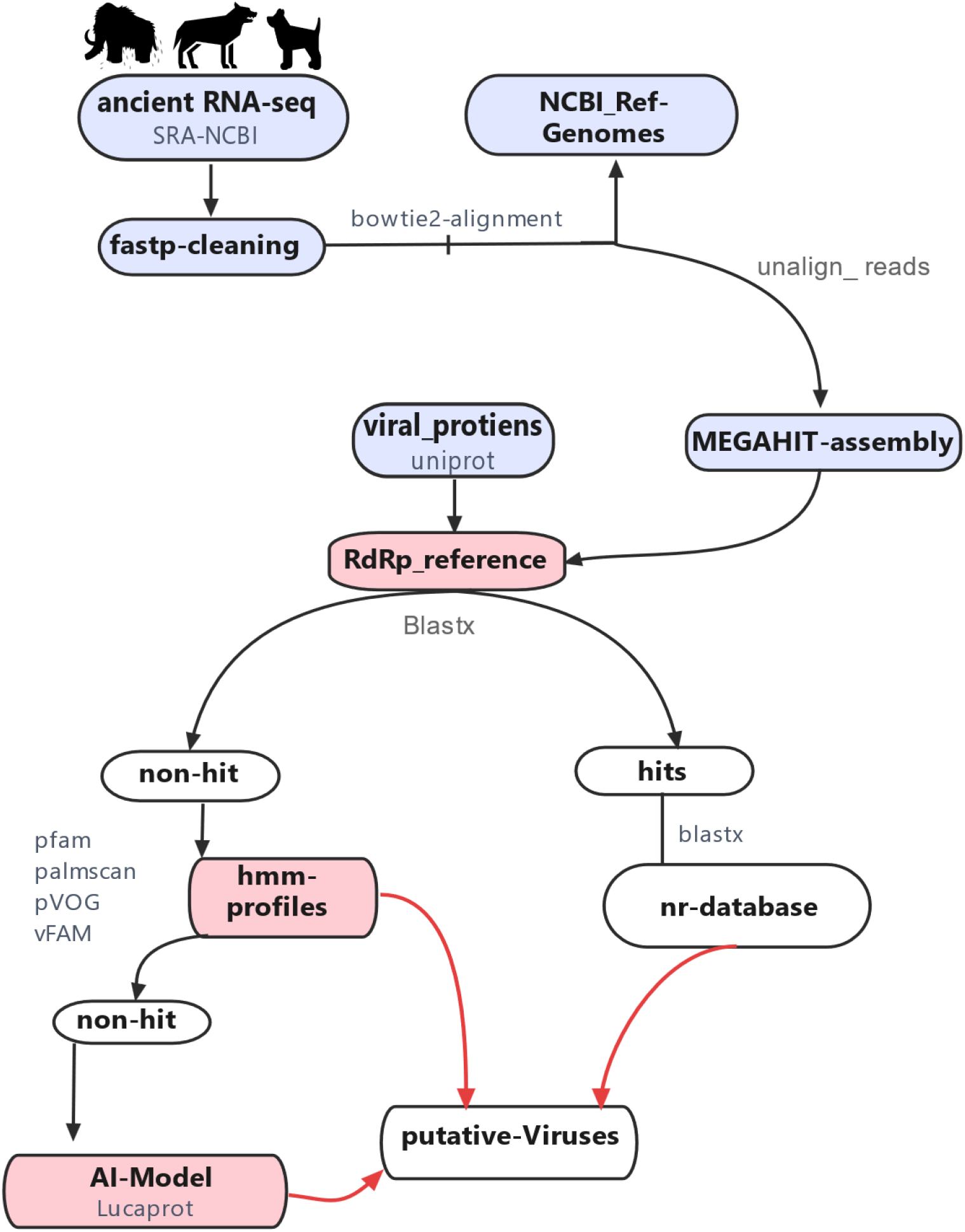
Overview of the viral-discovery pipeline. Raw RNA-seq reads from extinct-species transcriptomes (NCBI SRA) were quality-trimmed and host-mapped; unmapped reads were assembled de novo (MEGAHIT) and screened against a curated RdRp reference protein set (blastx). Hits were validated against the NCBI nr database, while non-hits were screened using HMM profiles (Pfam, PalmScan, pVOG, vFam), and the Lucaprot deep-learning model to recover additional divergent viral candidates. All positive hits were merged into a final set of putative viral contigs.

## Methods

### Data Acquisition and Pre-processing

Raw transcriptomic sequencing data (SRA) from ancient and extinct species were retrieved from the NCBI Sequence Read Archive (Supplementary Table 1). The targeted datasets included specimens from woolly mammoth (*Mammuthus primigenius*), Tasmanian tiger (*Thylacinus cynocephalus*), and gray wolf (*Canis lupus)* spanning four types of tissues: muscle, skin, liver, and cartilage (Figure 2). Fastp version 0.23.4 was used to trim adapter sequences and low-quality reads.

**Figure 2.**
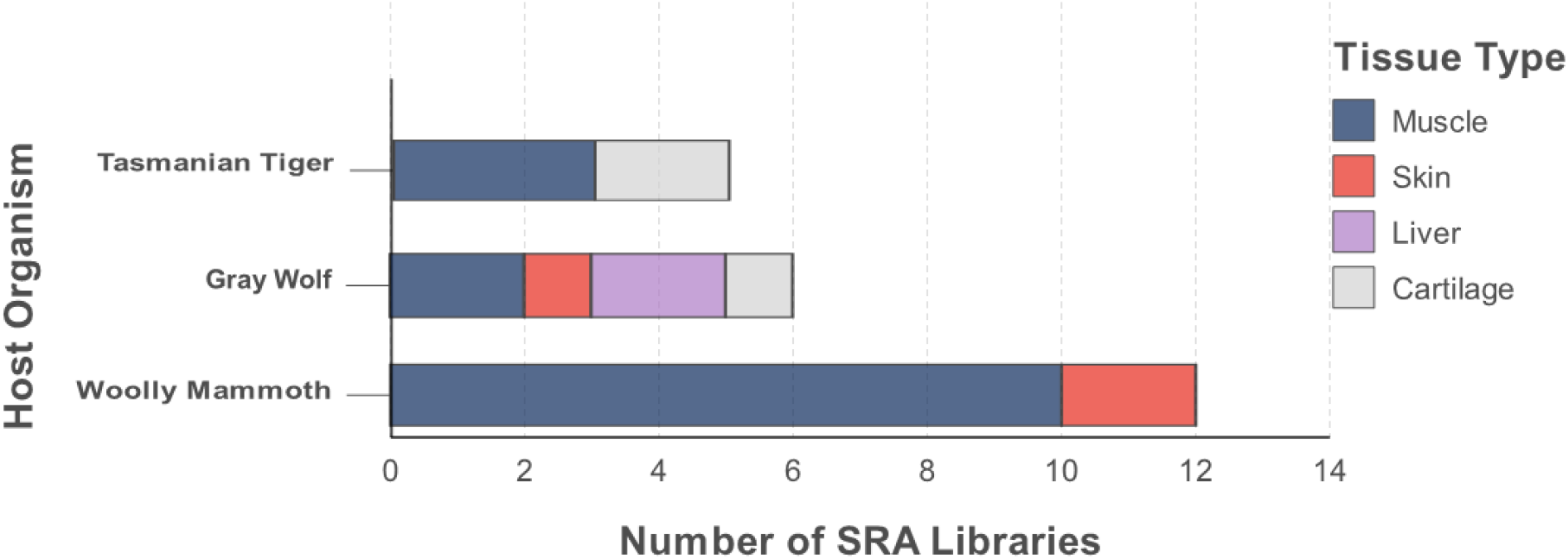
Distribution of SRA Libraries by Host Organism and Tissue Type.

### Host Removal

Quality-filtered reads were aligned with the reference genome from NCBI (*Canis familiaris, Loxodonta africana,* and *Sarcophilus harrisii (*Tasmanian devil) as proxies because of their good quality) using Bowtie2. Unaligned sequences were de novo assembled into contigs with MEGAHIT version 1.2.9 and the default metagenomic settings.

### Viral Sequence Identification and Annotation

Assembled contigs were directly screened against a custom viral protein database using Diamond BLASTx (e-value < 1e-5), as well as translated into predicted protein sequences using Prodigal’s-meta parameter (v2.6.3) (Hyatt, Chen et al. 2010). We extracted contigs with significant hits to viral hallmark genes, particularly the RNA-dependent RNA polymerase (RdRp). To ensure comprehensive detection, profile Hidden Markov Models (HMMs) for viral RdRp domains (Pfam and vFam) were applied using HMMER (Charon, Buchmann et al. 2022, Sakaguchi, Nakano et al. 2024). To improve sensitivity for highly divergent sequences, the deep-learning tool Lucaprot was additionally used to identify divergent RdRp-like contigs from non-hits. Candidate RdRps were then evaluated with HHblits (Remmert, Biegert et al. 2012) and PalmScan (Babaian and Edgar 2022) to assess completeness of the catalytic A, B, and C motifs; contigs lacking these motifs were considered incomplete or partial and were excluded from downstream phylogenetic analysis. Confirmed viral contigs were reassembled using CAP3 (-o 16). Putative species-level viral operational taxonomic units (vOTUs) were delineated using CD-HIT at an 80% threshold, with the conserved RdRp domain serving as the hallmark gene for RNA viruses. Because DNA viruses lack an RdRp hallmark gene, all DNA-virus-like sequences identified during screening were retained for downstream evaluation.

### Phylogenetic Analysis

The initial blastx results were used to determine the taxon of the putative viruses. To improve resolution, amino acid sequences of the putative RdRp domains from the novel viral contigs were aligned with reference RdRp sequences from the corresponding ICTV metadata using MAFFT v7.475 (L-INS-i strategy). Ambiguously aligned regions were trimmed using trimAl. We constructed maximum-likelihood phylogenetic trees using IQ-TREE2 with the best-fit substitution model selected by ModelFinder, and assessed branch support using ultrafast bootstrap approximation (UFBoot; 1000 replicates).

### Protein Structure Prediction and Structural Alignment

Since the novel viral RdRps exhibited low sequence identity (<55%) with known structures, we employed AlphaFold2 (Jumper, Evans et al. 2021) to predict the three-dimensional structures of the recovered proteins. The resulting models were evaluated based on pLDDT (predicted Local Distance Difference Test) confidence scores, and structural alignments between the novel viral RdRp models and their closest known relatives were performed using the TM-align algorithm in PyMOL. Conservation of the core polymerase domains, specifically motifs A, B, and C, which are responsible for substrate binding and catalysis, was assessed both visually and computationally.

### Annotation of Additional Viral Proteins

To identify and functionally annotate additional proteins encoded by the viral contigs beyond the RdRp, we employed complementary protein-domain search approaches. For open reading frames (ORFs) identified on the negative strand, we generated the reverse complement to represent the positive-sense coding strand for genomic organization analysis. These were then screened against the Pfam database using HMMER (v3.3) to detect conserved protein domains and families (Mistry, Chuguransky et al. 2021). In parallel, we searched the NCBI Conserved Domain Database (CDD) with RPS-BLAST (Reverse Position-Specific BLAST) to find conserved domains and functional motifs in the predicted viral proteins (Marchler-Bauer, Derbyshire et al 2015). Hits with an e-value less than 1e-5 were considered significant and saved for further investigation. This combined strategy allowed for the identification of potential auxiliary proteins, such as capsid components and movement proteins, that are typical of the viral families to which the novel contigs belong, providing additional evidence for their taxonomic placement and genomic organization.

## Results

Extensive mining of the ancient SRA libraries identified multiple contigs containing viral hallmark genes from samples SRR8090318 and SRR8090326, which are from an ancient wolf population. This fossil was found at the Syalakh site in northern Siberia, approximately 40 km from the village of Tumat. The geographic coordinates are 71.007500 N, 139.888889 E. This area is in the Ust-Yansky District of the Sakha Republic (Yakutia), in the Russian Federation. The samples were processed at the GLOBE Institute, University of Copenhagen. The sequencing itself was performed on an Illumina HiSeq 2500 platform.

**Figure 3:**
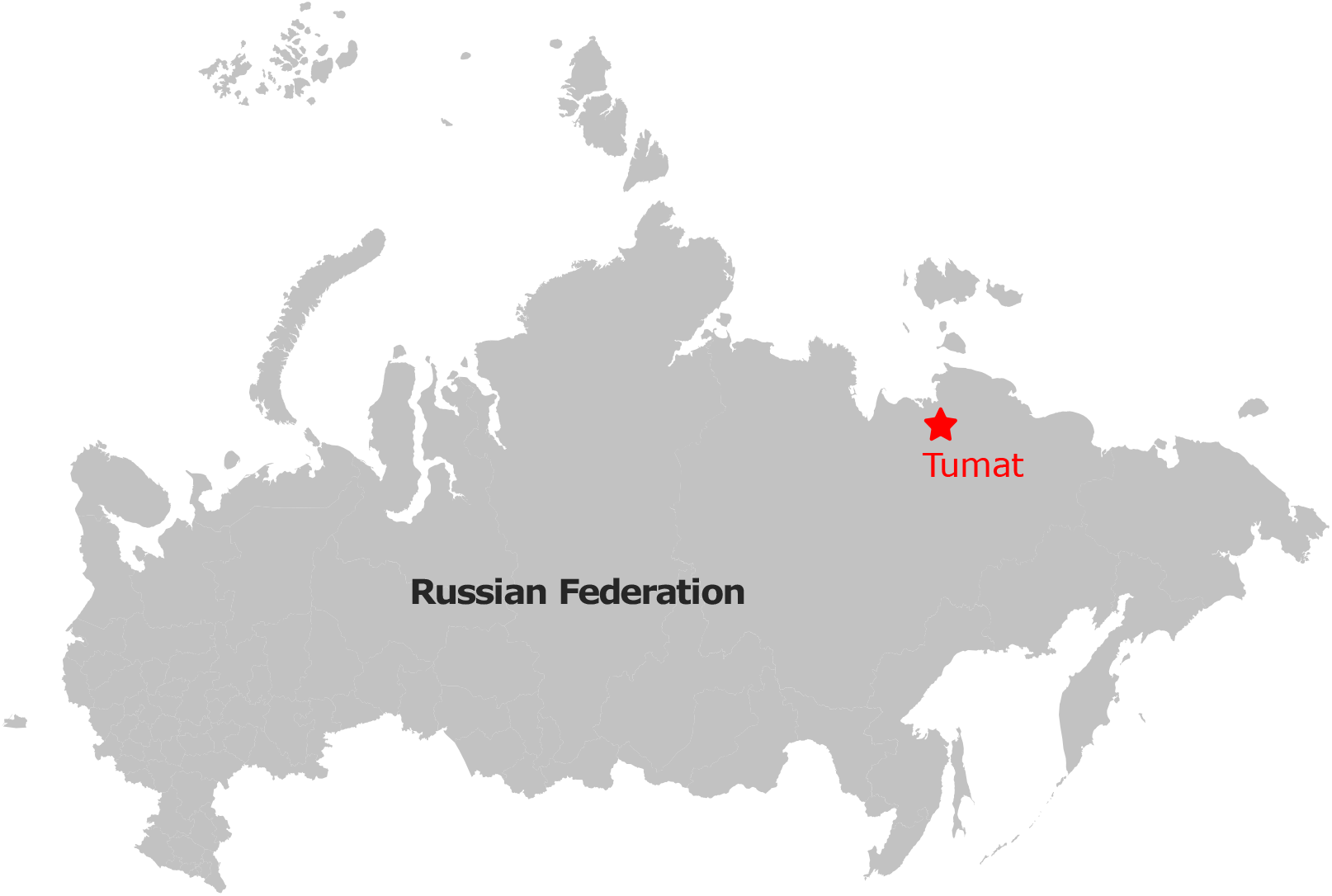
Location of the Tumat site (Sakha Republic, Russia). RNA-seq from a ∼14,300-year-old ancient *Canis lupus* specimen recovered at this permafrost locality yielded two novel viruses.

After stringent filtering, two distinct novel RNA viral sequences were retained for in-depth characterization. **Putative novel mitovirus** Contig1 (1,890 nt): Recovered from a 14,300-year-old Pleistocene *Canis lupus* library (SRR8090318 and SRR8090326), this contig encodes a 630-amino-acid (aa) putative RdRp. BLASTp and HMMER analyses identified its top hit as a *Rhizoctonia solani mitovirus* from the genus *Duamitovirus* with a low amino acid identity of approximately 57.6% and near-complete (∼98%) query coverage.

### Novel unclassified fungal virus Contig2 (1,566 nt)

Also recovered from the same libraries, this contig encodes an RdRp of approximately 496 aa. It shares 60.8% amino acid identity with *Rhizoctonia solani dsRNA virus 1* from the genus *Orthocurvulavirus*. The second genomic segment (822 nt) of this virus contains a single ORF encoding a putative protein of 279 amino acids. BLASTp and InterProScan analyses revealed no significant homology to known proteins or conserved functional domains, and therefore second segment was annotated as a hypothetical protein. This is consistent with the genomic organization of other *orthocurvulaviruses*, where the second segment often encodes a protein of unknown function.

The same sequencing runs also yielded a full genome of the *Escherichia* phage phiX174. A comparison with the reference phage genome (accession J02482) revealed a conserved genomic backbone, with capsid (F, G, H) and replication (A, B) genes sharing 100% amino acid identity. Read-degradation analysis verified that this sequence corresponded to the laboratory spike-in control commonly employed during Illumina sequencing, giving an internal benchmark for an undegraded, modern nucleic acid template against which the ancient viral contigs could be compared. No RNA viruses matching known mammalian pathogens were found in any of the ancient libraries, implying either the absence of active RNA viral infection at the time of death, destruction of mammalian RNA viruses over millennia, or loss of such transcripts during historical library preparation.

### Phylogenetic analysis and virus classification

We created maximum-likelihood phylogenetic trees using the RdRp amino acid sequences to assess the novel viruses’ evolutionary ties. Contig1(1890 nt) formed a discrete, well-supported branching clade within the genus *Duamitovirus*, clearly different from existing species in this genus (Figure 4). The second viral sequence Contig2 (1566 nt) clustered within the genus *Orthocurvulavirus* suggesting it represents a novel species within this genus, most closely related to a *Rhizoctonia solani dsRNA virus* (Figure 5).

**Figure 4.**
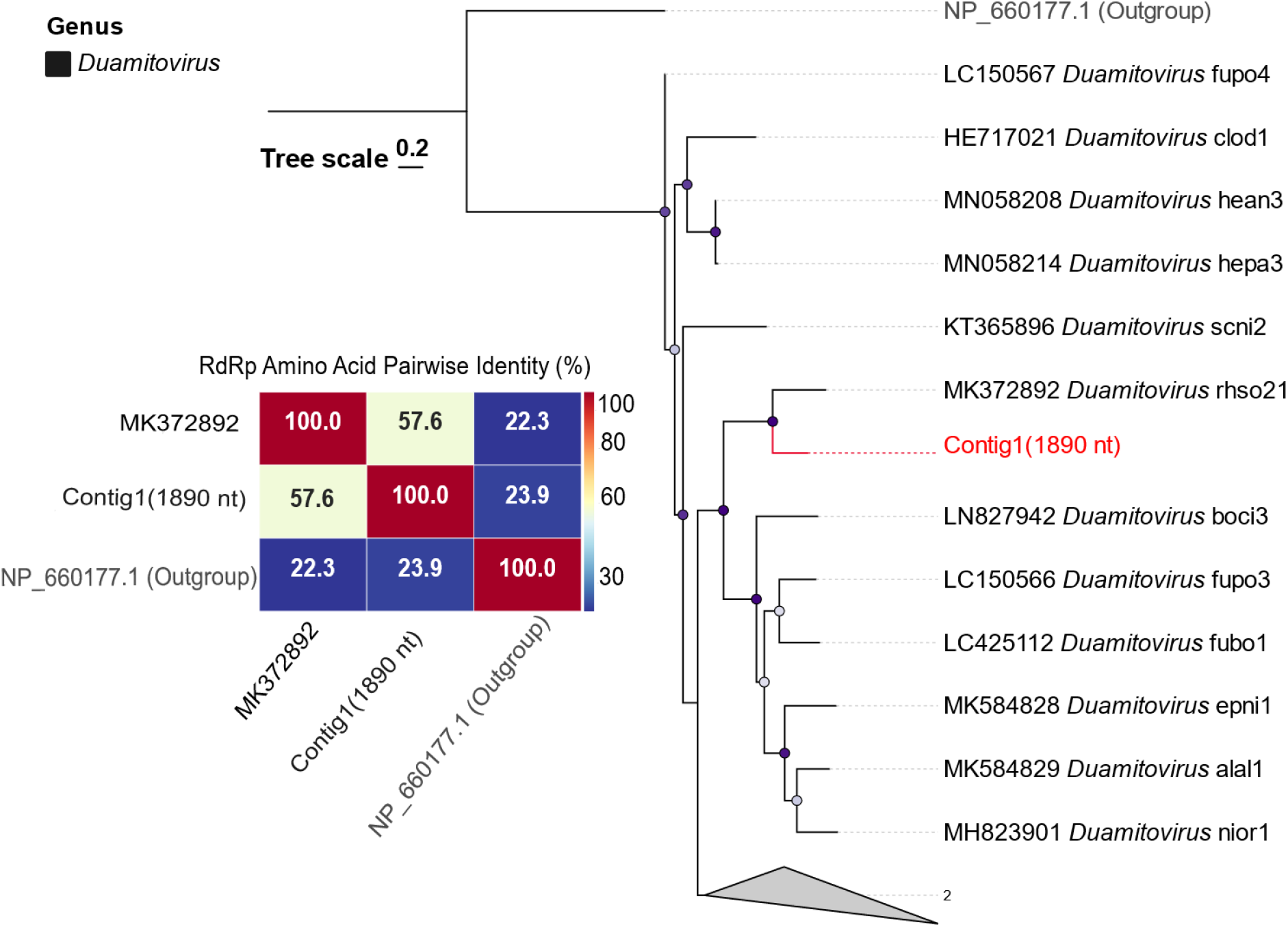
Phylogenetic placement and pairwise amino acid identity of the novel Contig1 RdRp sequence. Maximum-likelihood phylogenetic trees showing the placement of Contig1 (red) within its respective established genera, alongside pairwise percentage-identity matrices. It shares 57.6% RdRp amino acid identity with its closest relative (MK372892).

**Figure 5.**
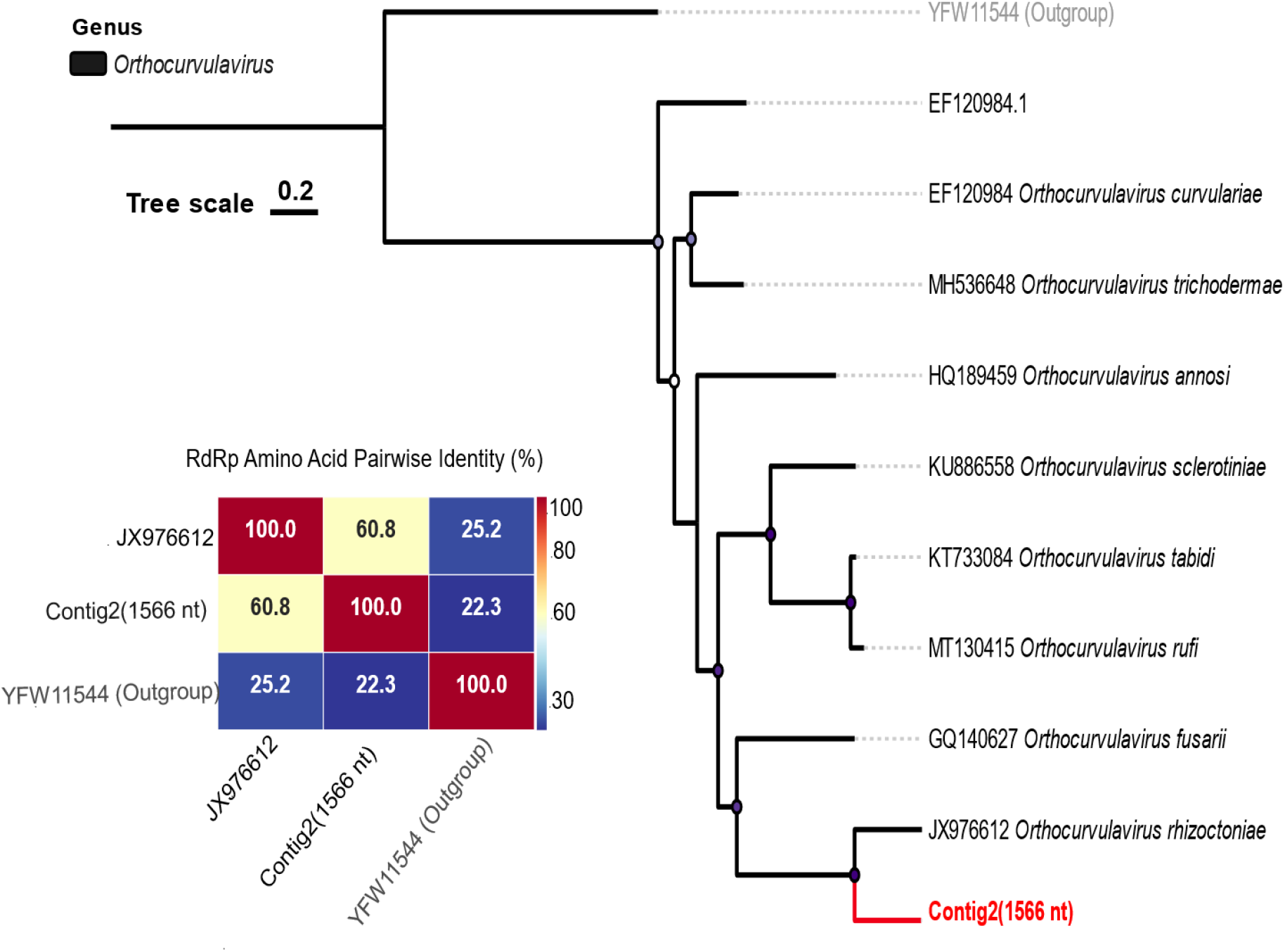
Phylogenetic placement and pairwise amino acid identity of the novel Contig2 (1566 nt) RdRp sequence. Maximum-likelihood phylogenetic trees showing the placement of Contig2 (red) within its established genus, *Orthocurvulavirus,* alongside pairwise percentage-identity matrices. It shares 60.8% RdRp amino acid identity with its closest relative (JX976612).

### Genomic organization and structural conservation of the RdRp core domains

For both novel viral entities, assembly from the ancient transcriptomes yielded at least a single, complete ORF encoding the RdRp. The genomic context, however, differs between the two contigs. Contig1 (1,890 nt) architecture, a solitary replicase gene, mirrors the simple monopartite organization of its closest known relative. In contrast, the closest relative of Contig2 has a bipartite genome comprising two segments, in which segment 1 encodes the RdRp protein and segment 2 encodes a multifunctional protein (involved in an unknown function). Because the recovered Contig2 (1566 nt) corresponded precisely to the RdRp-encoding segment 1, we infer that we captured the primary replicative module of this bipartite system and with additional exhaustive screening, another segment2 encoding ORF orthologous to the segment-2 protein of unknown function was confidently detected in these libraries and corresponds to a hypothetical protein of (*Rhizoctonia cerealis orthocurvulavirus*) with 65.95% identity and query coverage of 92% (Figure 6F). Because the RdRp sequences shared pairwise amino acid identities of ≤60% with their closest characterized homologs, substantially below the ICTV’s standard 80–90% species-demarcation threshold for these viruses, but their phylogenetic tree does not root them outside of established genera, we sought structural validation of their functionality. AlphaFold2 predictions yielded high-confidence models (mean pLDDT > 80) for all proteins, and especially structural alignment of RdRps with the nearest known relatives revealed a canonical viral right-handed polymerase fold. The catalytic core comprising the ABC domains, including the highly conserved motifs A (GDN), B (SG…T), and C (GDD), showed remarkable structural conservation despite the deep sequence divergence (Figure 6). This structural integrity indicates that the ancient identified contigs encode functional RdRps rather than degraded pseudogenes, and detectable read expression from these loci confirms active transcription within the ancient libraries; our sensitive pipeline detected positive as well as negative strands, indicating the active replication of these viruses. According to ICTV criteria, both sequences were classified as novel species within their established genera.

**Figure 6.**
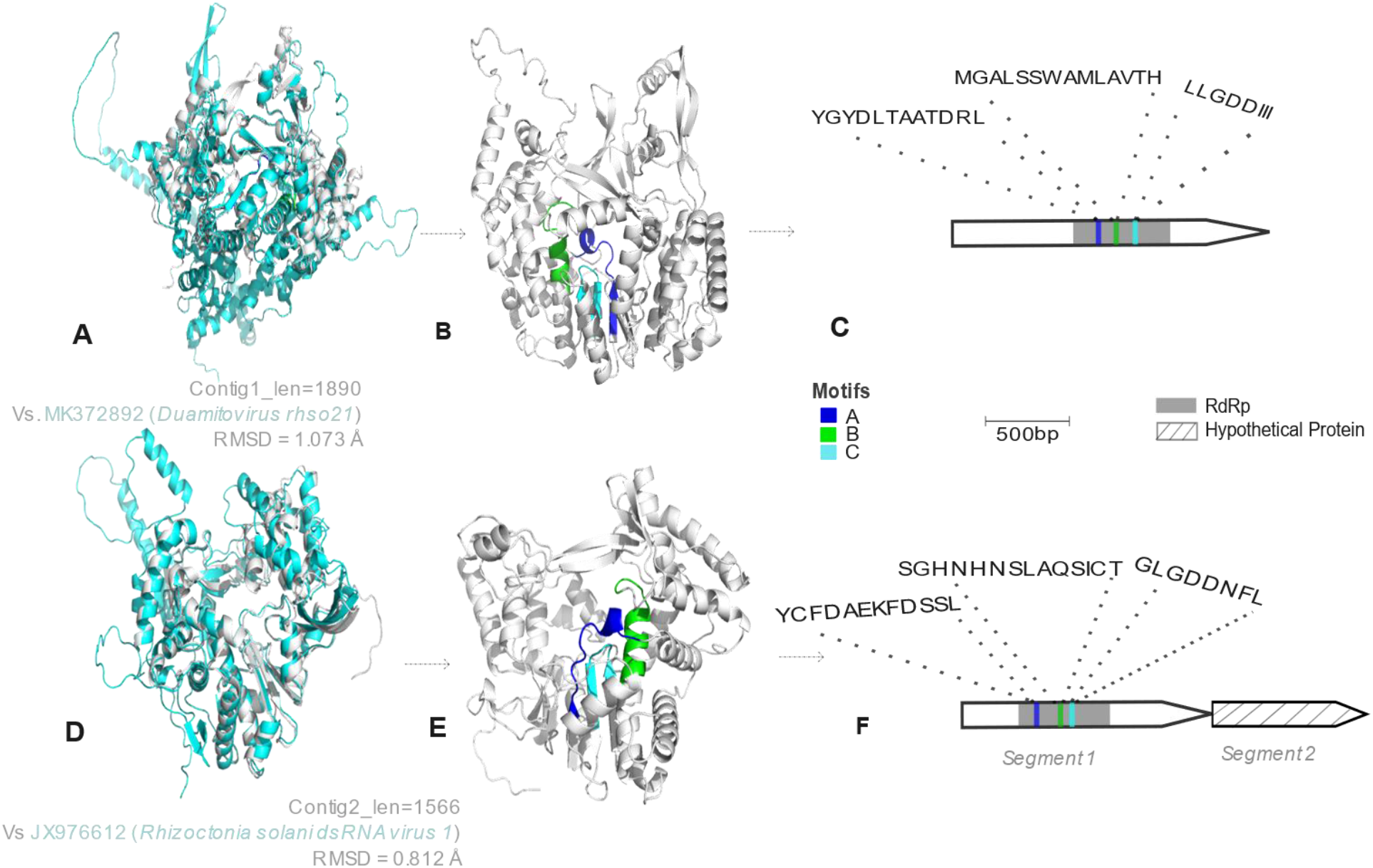
Structural prediction and genomic organization of the two novel ancient viruses. **(A)** AlphaFold2-predicted three-dimensional structures of Contig1 and its close relative MK372892 (*Duamitovirus rhso21*) were obtained for both proteins (mean pLDDT > 80). Structural alignment shows RMSD = 1.073 Å. **(B)** The conserved catalytic motifs (A, B, and C) within the polymerase fold. **(C)** Schematic representations of the recovered genomic Contig1 (1890 nt) shows that the sequence contains a single complete open reading frame encoding the RNA-dependent RNA polymerase with conserved catalytic motifs (A, B, and C) indicated; key residues for metal coordination (e.g., the GDD motif) are shown in sticks**. (D)** Contig2 RdRp shows an RMSD = 0.812 Å structural alignment with JX976612 **(E)** The conserved ABC domains within the RdRp-encoding gene **(F)** RdRp and segment2 (hypothetical protein) of a bipartite genome.

### Authentication of the novel viral sequences

To rigorously authenticate the ancient origin of the identified viral sequences and definitively exclude the possibility of modern laboratory or environmental contamination, we employed a dual-strategy validation framework combining comparative degradation profiling with exhaustive database-exclusivity analysis. First, we leveraged the unexpected co-assembly of a complete *Escherichia* phage phiX174 genome from the same sequencing libraries. While the phiX174 genome is a standard spike-in control for Illumina sequencing platforms, its presence in these libraries is consistent with its standard library preparation. This sequence exhibited 100% nucleotide identity to the reference phiX174 genome (accession J02482) and assembled as a fully circularized molecule, with all hallmark genes—including Microvir_H, Microvir_J, Phage_C, Phage_F, Phage_G, Phage_GPA, Protein_K, and gpD—achieving complete coverage (Figure 7).

**Figure 7.**
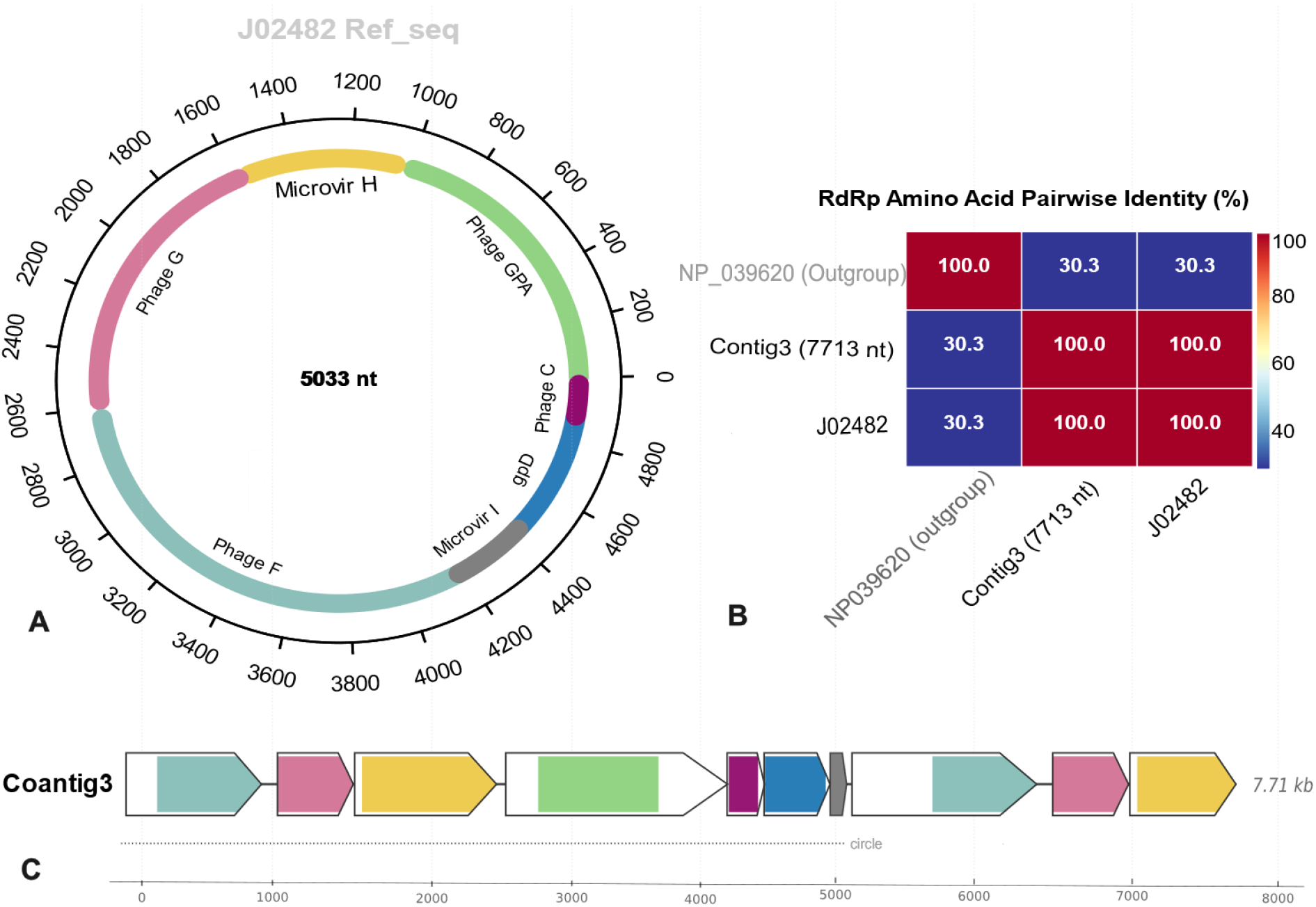
Assembly and annotation of the control phage sequence recovered from sequencing run SRR8090328. The assembled contig was mapped against the reference *Microviridae* J02482 (phiX174-related) genome. **(A)** Reference annotations, coding regions including Microvir_H, Microvir_J, Phage_C, Phage_F, Phage_G, Phage_GPA, Protein_K, and gpD. **(B)** The capsid protein shows 100% nucleotide identity to the reference with zero substitutions, confirming the presence of a laboratory-derived spike-in control. **(C)** Contig3 (7.71 kb) assembled from ancient SRA libraries.

The read profile displayed a uniformly intact length distribution and continuous coverage across the entire genome, consistent with a high-quality, non-fragmented nucleic-acid template. This phage sequence corresponds to the standard laboratory control routinely used for sequencing calibration, but here it may serve as an internal benchmark for a modern, intact template against which the degradation state of the ancient viral contigs could be objectively compared (Figure 8b).

**Figure 8:**
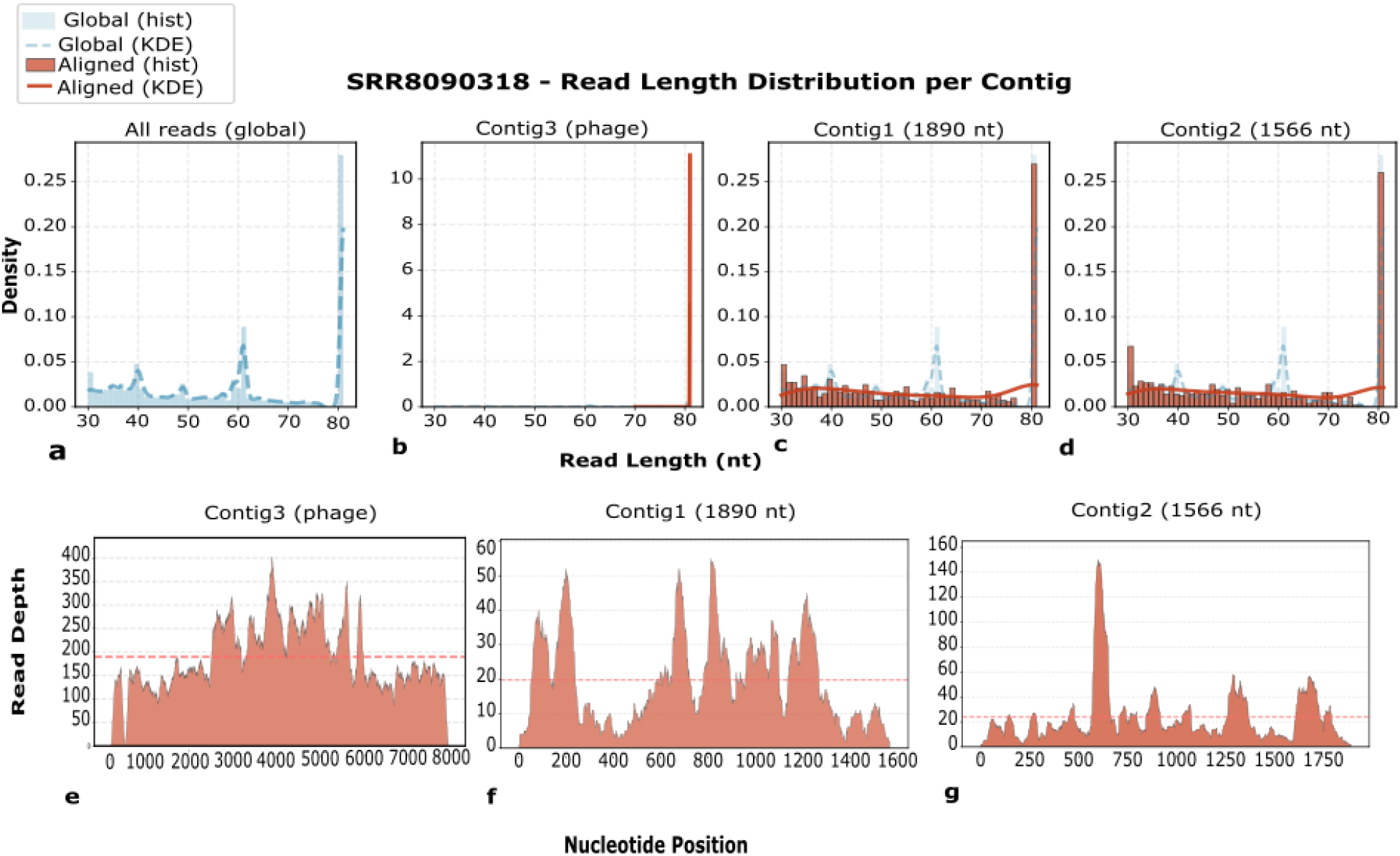
RNA-degradation characteristics of ancient viral contigs compared to a laboratory phage. **(A)** Charts illustrate the library’s global read-length distribution and coverage fragmentation**. (B)** Read-length distribution and coverage fragmentation charts for Contig3 (phage virus): The phage control has a constant, intact read-length distribution with minimal fragmentation. **(C**) Contig1(1890 nt) has severely fragmented read-length profiles and inconsistent coverage depths, indicating substantial postmortem RNA hydrolysis and chemical degradation over time. **(D)** Contig2 (1566 nt) has severely fragmented read-length profiles and inconsistent coverage depths, indicating substantial postmortem RNA hydrolysis and chemical degradation over time**. (E-G)** Nucleotide position coverage of raw reads for each viral contig. The distinction between perfect phage control and degraded ancient contigs shows that the revealed mycovirus sequences are real historical RNA molecules rather than laboratory contamination.

In stark contrast, the read profiles of both viral contigs (Contig1 and Contig2) were extremely fragmented, with short, discontinuous alignments and significantly varied coverage depths (Figure 8c-d). This pattern is indicative of substantial postmortem RNA hydrolysis caused by ribonucleolytic breakage over time. The consistent presence of these degradation signatures across both novel contigs, combined with their complete absence in the simultaneously processed phage control, clearly distinguishes the mycoviruses as genuine historical RNA molecules rather than recent laboratory contaminants.

Furthermore, to rigorously rule out the possibility of common environmental or fungal contaminants, we performed an exhaustive search against the Logan planetary-scale SRA database, which comprises approximately 4.93 million transcriptomes and 0.13 million metatranscriptomes. Across a range of coverage criteria (0.3-0.8), hits for both Contig1 and Contig2 were obtained from the sole ancient *Canis lupus* library pair (SRR8090318 and SRR8090326), with no detections in any other metatranscriptomic or transcriptomic dataset (Figure 9). One exception was Contig1, which produced a single hit in library SRR8090319 at the lowest coverage criterion (0.3). This library is derived from the same study and organism as the two primary source libraries were, and no contig was obtained during de novo assembly in this library, implying that the relevant viral fragment was not present at sufficient depth or continuity for assembly. Rather than signaling a false positive, this low-coverage detection most likely represents a short, unassembled sequence fragment from the same underlying viral population, demonstrating the sensitivity of our read-level search technique in comparison to assembly-dependent detection alone. This significant database rarity categorically rejects the appearance of these sequences as widespread modern fungal contaminants, which would otherwise be expected to appear across multiple environmental or mycological datasets.

**Figure 9.**
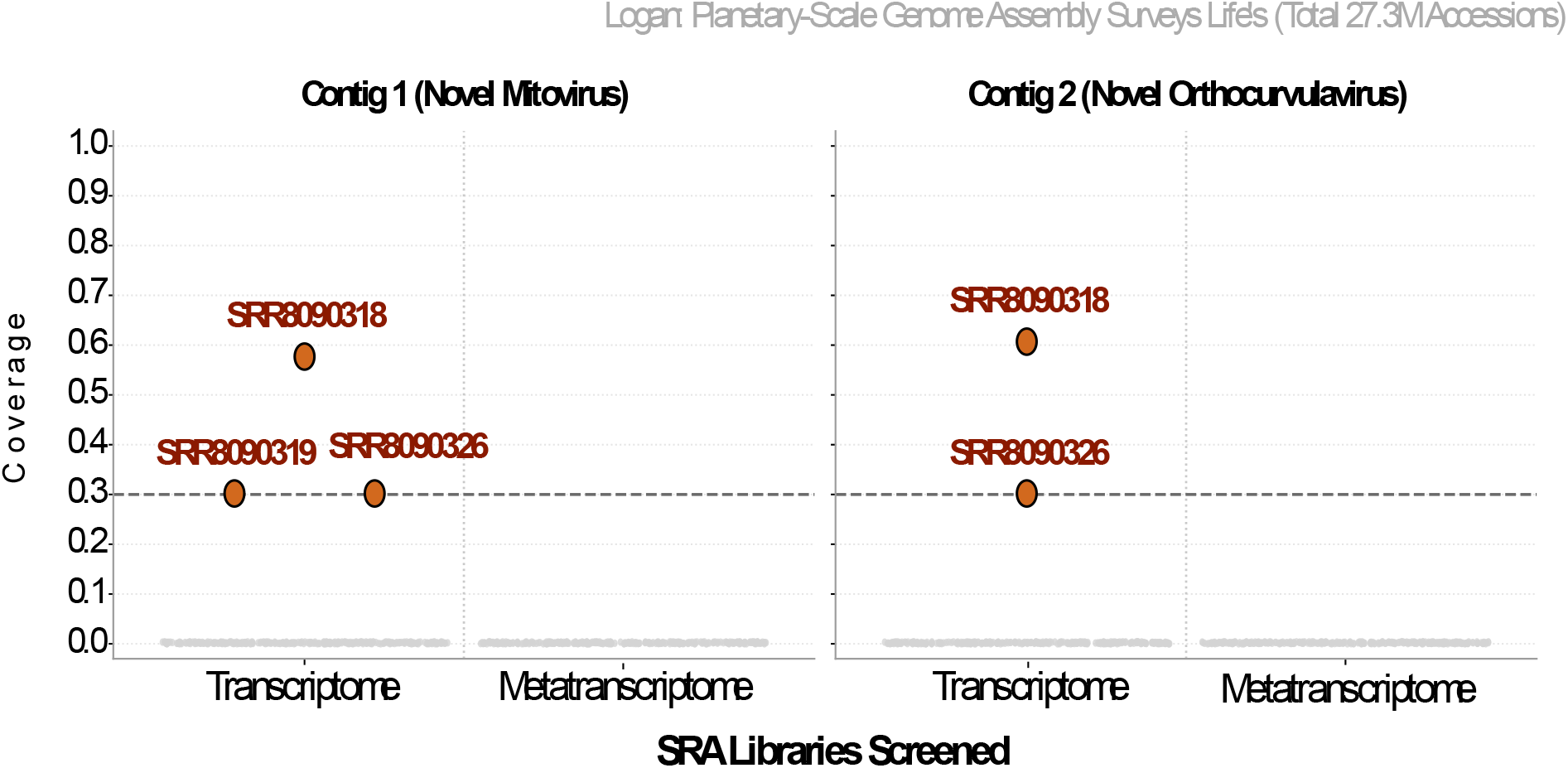
Logan-based global screening across 27.3 million accessions reveals the occurrence of two novel RNA viruses. Contig 1 (novel *Duamitovirus*) was detected solely in SRR8090318, SRR8090319, and SRR8090326, while Contig 2 (novel *Orthocurvulavirus*) was restricted to SRR8090318 and SRR8090326. No homologous sequences were found in any other publicly available data, confirming that these viruses are unique to the ancient Tumat wolf carcass at the time of death and currently have not been sampled elsewhere in the biosphere.

Collectively, these complementary lines of evidence the degraded RNA fragmentation profiles, the pristine integrity of the internal phage calibrator, and the absolute database exclusivity of the contigs robustly authenticate the ancient provenance of the novel mycoviruses. We conclude that these sequences represent genuine historical viral entities preserved within the ancient transcriptome, reflecting the presence of a fungal microbiome associated with the extinct canid specimen at the time of death.

## Discussion

We identified two novel RNA viruses exclusively associated with this extinct fauna. We have named the first virus *Duamitovirus tumati* (Contig 1), which belongs to the family *Mitoviridae* and is biologically expected to encode only the RNA-dependent RNA polymerase (RdRp), as mitoviruses are capsidless “naked RNA” viruses that lack structural proteins. We retrieved a nearly complete genome for this unique virus, with the RdRp serving as the sole coding region. The second virus, termed *Orthocurvulavirus tumati dsRNA* (Contig 2), belongs to the *Curvulaviridae* family and has a bipartite genome organization consisting of two segments. Segment 1 encodes the RdRp, and Segment 2 encodes a putative protein with uncertain function. Despite comprehensive structure prediction using AlphaFold and sensitive homology-based searches against publicly available databases, we could not assign a clear function to this protein. A weak similarity to *Togovirus* capsid proteins was found, but the matching root-mean-square deviation (RMSD) values were insufficient to provide a reliable structural or functional interpretation. Transmembrane topology and subcellular localization prediction with Phobius revealed a substantial non-cytoplasmic domain spanning residues 1-279, implying that this protein is membrane-anchored, secreted, or confined to an extra-cytoplasmic compartment. However, its particular biological function remains unknown. Collectively, our data show that both viruses are unique species within their respective genera, adding to the growing diversity of RNA viruses found in ancient microbiomes. Over millennial periods, RNA is gradually hydrolyzed due to the intrinsic lability of the ribose 2′-hydroxyl group, which promotes transesterification and backbone cleavage. Preservation requires exceptional conditions—such as rapid desiccation, continuous subzero temperatures, or chemical cross-linking—that inhibit nucleophilic attack on the phosphodiester backbone (Li and Breaker 1999). The recovery of ancient complete RNA sequences, both positive and negative strands, from the canid specimen in this study supports the action of one or more such preservation mechanisms (Anderson 2025). During de novo assembly from fragmented, damaged reads, the most conserved and structurally constrained genomic regions are preferentially recovered. The RdRp is the most evolutionarily conserved gene across RNA virus lineages, owing to strong structural and functional constraints on the catalytic motifs required for replication. Consequently, RdRp fragments are more likely than the more rapidly evolving structural genes to retain sufficient sequence identity for BLAST-based recognition and to assemble into contiguous sequences. This mechanistic explanation accounts for the recovery of complete RdRp ORFs.

The application of metatranscriptomics to publicly archived sequencing datasets has accelerated the discovery of novel viruses. In line with this approach, we extended such mining to transcriptomic libraries derived from extinct and ancient species. Prior work on modern host transcriptomes, for example analyses spanning over 17,000 rice transcriptomes (Zhu, Raza et al. 2025), has revealed extensive RNA viral diversity. This study, mining ancient mammalian libraries, recovered fungal-associated viral sequences. The absence of detectable mammalian RNA viruses in these specimens is likely multifactorial. First, the sampled individuals may simply not have been experiencing an active, viremic RNA viral infection at the time of death, a stochastic factor that cannot be excluded. Second, the inherent chemical instability of RNA may render mammalian viral genomes particularly susceptible to rapid hydrolysis, potentially degrading them beyond the limits of assembly even under favorable preservation conditions. In addition, historical sequencing of extinct megafauna, including the Tasmanian tiger and woolly mammoth, was predominantly optimized for genomic DNA recovery or enriched for host nuclear regions, leaving limited sequencing depth for incidental RNA viral transcript capture.

Despite limitations, the discovery of two novel RNA viruses that share 57.6% and 60.8% RdRp amino acid identity with their closest known relatives, placing them in the *Mitoviridae* and *Curvulaviridae* families and a related fungal viral lineage, represents a significant contribution to viral paleodiversity. The presence of *Mitoviruses* in ancient specimens shows that fungal colonizers were present during death or early diagenesis. Fungal colonization of carcasses is well established, and these fungi appear to have harbored chronic viral infections. Multiple lines of evidence support the classification of these genomes as novel species. The International Committee on Taxonomy of Viruses (ICTV) has traditionally employed an 80-90% threshold for species delineation, but RdRp amino acid identity is significantly lower in these species. Furthermore, AlphaFold-based structural predictions provide additional evidence for viral functionality: despite tens of thousands of years of sequence divergence, the polymerase’s conserved catalytic ABC motifs and tertiary architecture remain mostly intact, consistent with the structural constraints imposed by viral replication. Beyond the discovery of two novel viruses, this study demonstrates that current viral discovery tools particularly AI-based approaches and structural prediction via AlphaFold have matured to the point where the primary limitation is no longer computational detection, but rather the physical recovery of ancient RNA (aRNA) from preserved specimens. Our results highlight that the field is now well-positioned to address long-standing questions about viral evolution, provided that improvements in aRNA extraction and preservation continue to advance.

### Limitations of this study

- Viral authentication of ancient origin was based on the degradation pattern, modern contamination assessment, and absence in the planetary-scale databases. We were unable to perform direct biochemical dating or deamination analysis.
- Despite structural prediction, segment-2 of Contig2 (1566 nt) remains a hypothetical protein. We were unable to predict the function with significant confidence.
- No experimental validation: RdRp functionality was inferred from AlphaFold2 models and motif prediction; no biochemical/enzymatic assays were performed.

In conclusion, this study shows that paleotranscriptomic mining of extinct-species databases is a promising method for identifying novel exogenous viruses. While mammalian ancient RNA viruses remain elusive, the discovery of these deeply divergent viruses broadens our understanding of the RNA virosphere over geological timescales and provides a methodological foundation for future paleovirological research aimed at reconstructing the ancient viral ecosystems associated with Earth’s extinct megafauna.

## Author Contributions

S.Z. conceived and designed the study. S.Z. performed the computational metatranscriptomic analyses, homology searches, structural predictions, degradation profiling, and drafted the original manuscript. Q.W. supervised the project, provided funding and resources, and reviewed and edited the manuscript. Both authors read and approved the final version.

## Acknowledgements

This work was supported by the Strategic Priority Research Program of the Chinese Academy of Sciences (Grant No. XDB0490000) and the Secretariat of the Alliance of National and International Science Organizations (CAS-ANSO). We thank the University of Science and Technology of China (USTC) Supercomputing Center for computational infrastructure.

## Data and Code Availability Statement

RNA-seq datasets used in this study were downloaded from a public repository accessible via the NCBI Sequence Read Archive (SRA) under the accession numbers provided in Supplementary Table 1. The consensus sequences of the viral candidates identified in this work have been deposited in NCBI GenBank under the submission identifier SUB16436308. Python, bash scripts, and viral sequence files are available on the GitHub repository: https://github.com/Syed-Zaheer-ud-Din/Noval-RNA-Viruses-Extinct-species-.git)

## Supplementary Material

**Table 1.** SRA RNA-seq libraries used in this study.

| SRA Accession | Host Organism | Sample / Tissue |
| --- | --- | --- |
| SRR31502636 | *Mammuthus primigenius* (woolly mammoth) | skin |
| SRR31502638 | *Mammuthus primigenius* (woolly mammoth) | skin |
| SRR31502639 | *Mammuthus primigenius* (woolly mammoth) | Muscle |
| SRR31502640 | *Mammuthus primigenius* (woolly mammoth) | Muscle |
| SRR31502641 | *Mammuthus primigenius* (woolly mammoth) | Muscle |
| SRR31502642 | *Mammuthus primigenius* (woolly mammoth) | Muscle |
| SRR31502643 | *Mammuthus primigenius* (woolly mammoth) | Muscle |
| SRR31502644 | *Mammuthus primigenius* (woolly mammoth) | Muscle |
| SRR31502645 | *Mammuthus primigenius* (woolly mammoth) | Muscle |
| SRR31502646 | *Mammuthus primigenius* (woolly mammoth) | Muscle |
| SRR31502647 | *Mammuthus primigenius* (woolly mammoth) | Muscle |
| SRR33328226 | *Mammuthus primigenius* (woolly mammoth) | Muscle |
| SRR8090318 | *Canis lupus* (gray wolf) | Muscle |
| SRR8090319 | *Canis lupus* (gray wolf) | Liver |
| SRR8090326 | *Canis lupus* (gray wolf) | Muscle |
| SRR8090324 | *Canis lupus* (gray wolf) | Skin |
| SRR8090327 | *Canis lupus* (gray wolf) | Liver |
| SRR8090328 | *Canis lupus* (gray wolf) | Cartilage |
| SRR23147616 | *Thylacinus cynocephalus* (Tasmanian tiger) | Muscle |
| SRR23147615 | *Thylacinus cynocephalus* (Tasmanian tiger) | Muscle |
| SRR23147614 | *Thylacinus cynocephalus* (Tasmanian tiger) | Muscle |
| SRR23147613 | *Thylacinus cynocephalus* (Tasmanian tiger) | Skin |
| SRR23147611 | *Thylacinus cynocephalus* (Tasmanian tiger) | Skin |

